# Innate immune recognition of a bacterial MAMP leads to conditional activation of pro- or anti-inflammatory responses

**DOI:** 10.1101/717256

**Authors:** Joo-Hong Park, Steve Cornick, Giulia Nigro, Gwladys Sevrin, François Déjardin, Ron Smits, Marion Bérard, Francina Langa, Ivo G. Boneca, Andrew Gewirtz, Benoit Chassaing, Nicolas Barnich, Philippe Sansonetti, Gérard Eberl

**Author notes:** These authors contributed equally to this work (Joo-Hong Park, Steve Cornick). School of Biological Sciences, Seoul National University, Seoul 08826, Korea (Joo-Hong Park).

## Abstract

Microbe-associated molecular patterns (MAMPs) are recognized by pattern recognition receptors (PRRs) of the innate immune system. Flagellin, the primary component of bacterial flagella, is recognized by membrane TLR5 and cytoplasmic NLRC4 receptors, which promote a vigorous pro-inflammatory response typically associated with bacterial infection. However, herein, we report that the nature of the flagellin-induced response is highly dependent on the physiological state of the tissue. Specifically, in the steady state, epithelial cell detection of flagellin orchestrates an anti-inflammatory response mediated by IL-33-dependent type 2 regulatory T cells while, in the context of injury, it induces a pro-inflammatory response mediated by myeloid cells, IL-18 and Th17 cells. Likewise, in the absence of infection, bacterial symbionts expressing high levels of flagellin induce a type 2 response. These data demonstrate that, depending on the inflammatory state of the milieu, MAMPs can function both as immunogens or tolerogens.

---

MAMPs, or pathogen-associated molecular patterns (PAMPs), were originally proposed to be recognized by PRRs on dendritic cells (DCs) in order for the immune system to distinguish self from non-self, and thereby trigger an inflammatory response against pathogens *(1)*. This concept was later modified to include danger-associated molecular patterns, or DAMPs, which signal tissue injury as a consequence of infection and trauma *(2)*. Both these views assume that the immune system responds to MAMPs and DAMPs through inflammation to eliminate microbes and injury. However, the recognition of MAMPs can also lead the development of the immune system and symbiosis with microbes. For example, upon recognition by NOD1, peptidoglycans released by the bacterial cell wall induce the formation of hundreds of intestinal lymphoid tissues involved in homeostasis of the host with microbiota *(3)*, and bacterial polysaccharides activate TLR2 and induce the generation of regulatory T cells (Tregs) that prevent inflammation and the rejection of the intestinal symbiont *(4)*.

In order to assess the impact of a single MAMP on the immune system in the absence of confounding tissue injury provoked by bacterial infection or MAMP injection, we generated a transgenic mouse system that allows for controlled spatial and temporal expression of bacterial flagellin. The *fliC* coding sequence derived from *Salmonella typhimurium* was codon-optimized for the mouse genome and inserted into a neutral bacterial artificial chromosome (BAC) devoid of genes, under control of a doxycycline (dox)-inducible *tetO*_*7*_ promoter (Fig. 1A). These mice, expressing a cytoplasmic form of flagellin, were crossed to *Villin*^rtTA^ mice that express the reverse tet transactivator (rtTA) under control of the *Villin* promoter *(5)*. Hence, when given dox, the *Villin*^rtTA^ x c*fliC* mice expressed flagellin in intestinal epithelial cells, as reported by the expression of the *FliC* transcript and EGFP (Fig. 1B), modelling the presence of bacteria at the mucosal barrier. Even though Villin is expressed in all intestinal epithelial cells *(5)*, only a fraction of epithelial cells expressed EGFP, possibly reporting only high expressors of flagellin. EGFP expression was not associated with a particular epithelial cell subset (data not shown), and no alteration in apoptosis nor proliferation was detected upon flagellin expression (fig. S1).

**Fig. 1.**
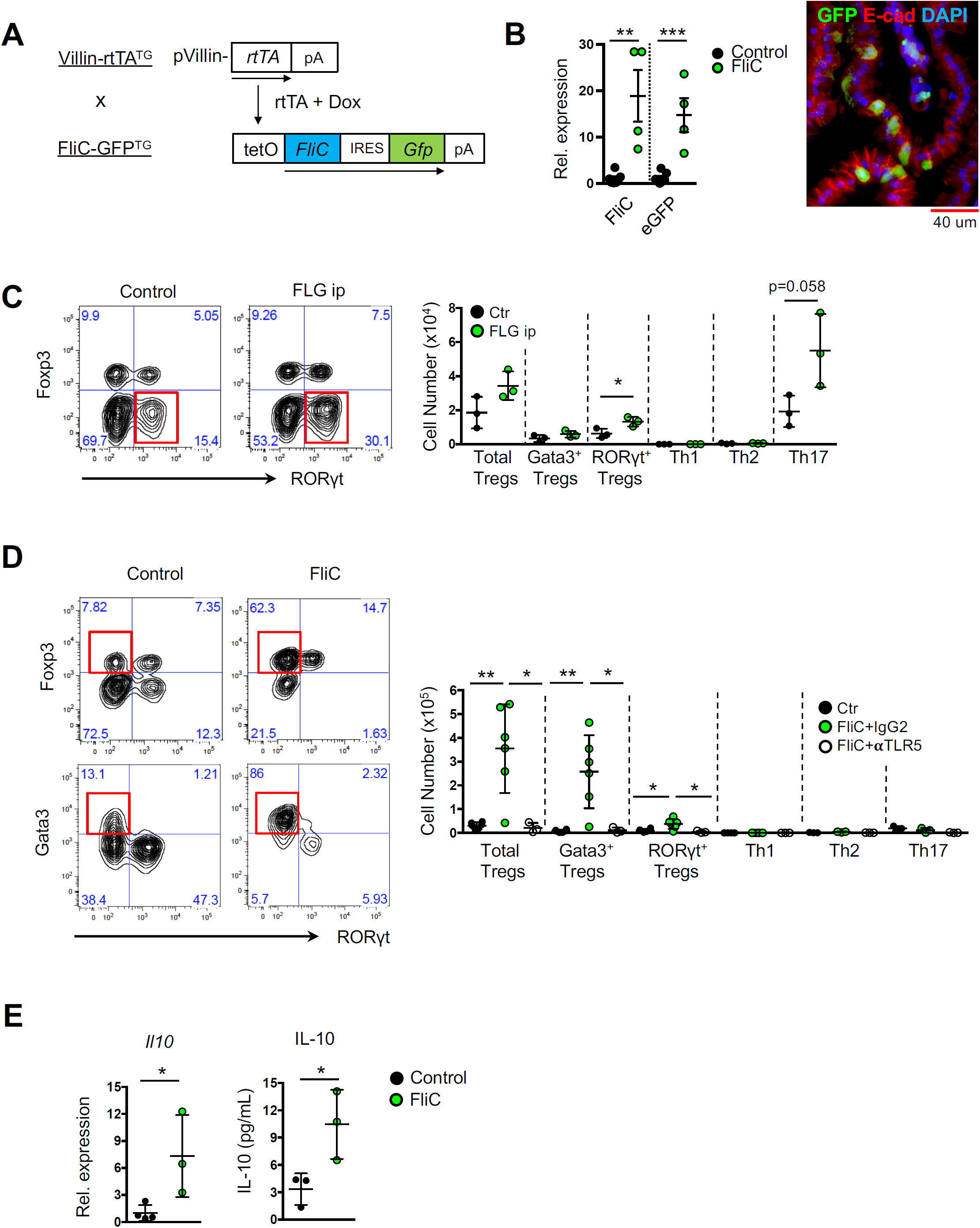
Flagellin induces an anti-inflammatory response at the steady state. (**A**) Schematic representation of the genetic strategy of *Villin*^rtTA^ x c*fliC* mice. The *fliC* coding sequence, followed by an *Ires-Gfp* cassette, was inserted into a bacterial artificial chromosome (BAC) as a vector for transgenesis. (**B**) Expression of transcripts for FliC and EGFP in small intestine epithelial cells (IECs). Immunofluorescence staining for GFP, E-cadherin and DAPI in IECs 3 days after doxycycline (dox) treatment in the drinking water. (**C**) Expression of Foxp3 and ROR*γ*t by T cells from small intestine lamina propria of C57BL/6 mice treated with PBS or recombinant flagellin intraperitoneally daily for 6 days (left). Cell numbers of Tregs and T helper (Th) cells in the small intestine (right). (**D**) Expression of Foxp3, Gata3 and ROR*γ*t by T cells from small intestine lamina propria of littermate controls and *Villin*^rtTA^ x c*fliC* mice treated with dox for 6 days (left). Cell numbers of Tregs and Th cells in littermate control and *Villin*^rtTA^ x c*fliC* mice treated with either isotype or anti-TLR5 neutralizing antibody (right). (**E**) Expression of *Il10* transcripts in the ileum (left), and production of IL-10 by T cells isolated from small intestines (right) of littermate control and *Villin*^rtTA^ x c*fliC* mice treated with dox. Error bars, s.d., *P<0.05, **P<0.01, ***P<0.001, as calculated by Student’s *t*-test.

Intravenous administration of recombinant flagellin induces an IL-23-mediated type 3 immune response dominated by IL-22 *(6, 7)*. In accordance with these results, we find that intraperitoneal administration of flagellin induces an expansion of Th17 cells and type 3 innate lymphoid cells (ILC3s), as well as of type 3 regulatory T cells (Tregs) that express the nuclear hormone receptor and transcription factor ROR*γ*t (Fig. 1C, fig. S2A). In contrast, when flagellin expression was induced in *Villin*^rtTA^ x c*fliC* mice, a vigorous anti-inflammatory response was generated in the small intestine through the expansion of type 2 Tregs producing IL-10, as well as a comparatively marginal expansion of Th2 cells and ILC2s, which express the transcription factor Gata3 (Fig. 1D and E, fig. S2B and S2C). Of note, type 2 Tregs are associated with repair responses in the intestine, muscle and adipose tissue *(8-10)*. Similar results were obtained using *Villin*^rtTA^ x s*fliC* mice that expressed a secreted form of flagellin (fig. S2D).

Retinoic acid (RA) is an efficient cofactor in the generation of Tregs *(11, 12)*, and the type 2 inducer cytokine IL-33 promotes the stability and function of type 2 Tregs *(8)*. Accordingly, RA was necessary for flagellin-induced expansion of Tregs (Fig. 2A, fig. S3A), and the expression of ST2, one of the two chains forming the receptor for IL-33, was increased in flagellin-induced Tregs (Fig. 2B). Furthermore, the frequency of type 2 Tregs, as well as their level of Gata3 expression, was decreased in the presence of neutralizing antibodies to IL-33 (Fig. 2B, fig. S3B). IL-33 is a nuclear protein that is released by epithelial and endothelial cells *(13)*. The intestinal epithelium of dox-treated *Villin*^rtTA^ x c*fliC* mice increased transcript expression for IL-33, as well as for the other type 2 inducer cytokines IL-25 and TSLP (Fig. 2C). It also increased expression of *Aldh1a2* coding for retinaldehyde dehydrogenase 2 (RALDH2) that converts dietary vitamin A into RA, and transcripts for TGF*β*1, both involved in the generation of Tregs *(11)*, indicating that flagellin expression by intestinal epithelial cells induces epithelial cells to express several factors promoting the generation of type 2 Tregs. Of note, *Aldh1a2* is also expressed by CD103^+^ dendritic cells (Fig. 2D), as previously reported *(14)*.

**Fig. 2.**
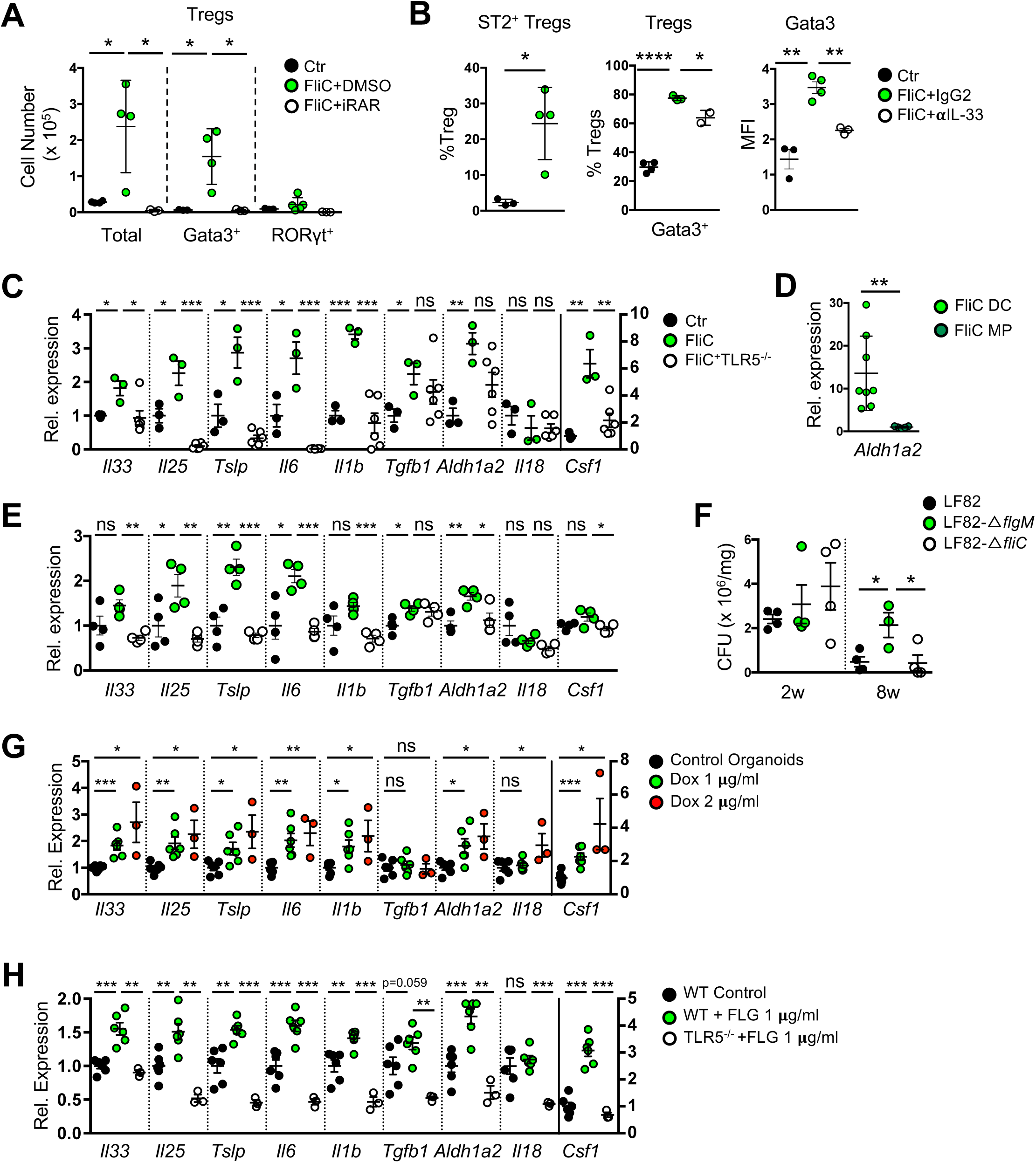
Flagellin-activated epithelial cells orchestrate an anti-inflammatory response. (**A**) Cell numbers of Tregs in small intestine lamina propria of control and *Villin*^rtTA^ x c*fliC* mice treated with dox and either DMSO or retinoic acid receptor (RAR) inhibitor intraperitoneally daily for 6 days. (**B**) Frequencies of ST2^+^ Tregs in dox-treated control and *Villin*^rtTA^ x c*fliC* mice, and frequencies of Gata3^+^ Tregs and MFI of Gata3 in Tregs in control and *Villin*^rtTA^ x c*fliC* mice treated with dox and either isotype or anti-IL-33 neutralizing antibody. (**C** and **D**) Gene expression in IECs of the ileum of dox-treated control and *Villin*^rtTA^ x c*fliC* mice, and expression of *Aldh1a2* transcripts in dendritic cells and macrophages from dox-treated *Villin*^rtTA^ x c*fliC* mice. (**E**) Gene expression in IECs of the ileum of germ-free mice colonized with wild-type, *ΔflgM* or *ΔfliC E.coli* LF82, 2 weeks after inoculation. (**F**) Numbers of colony forming unit (CFU) in feces of LF82-colonized germ-free mice. (**G**) Gene expression in intestinal organoids derived from *Villin*^rtTA^ x c*fliC* mice. Organoids were treated with doxycycline 2 days after the initiation of the cultures of crypt stem cell and analysed by qPCR 5 days after induction. (**H**) Gene expression in intestinal organoids from wild-type or TLR5 KO mice, treated with recombinant flagellin 2 days after the initiation of the cultures and analysed as in (G). Error bars show ± SEM, ns means not significant, *P<0.05, **P<0.01, ***P<0.001, as calculated by Student’s *t*-test.

In order to assess whether transgenic expression of flagellin in intestinal epithelial cells functionally reflected flagellin produced by intestinal bacteria, we mono-colonized germfree mice with a series of *Escherichia coli* strains. Mono-colonization of germfree mice with a *ΔflgM* mutant of the *E. coli* strain LF82 *(15)*, which secretes high levels of flagellin, induced a similar type 2 expression pattern in epithelial cells, whereas the parental strain, or a *ΔfliC* mutant that does not express flagellin, failed to do so (Fig. 2E). Even though Tregs did not significantly expand in this context, the expansion of Th17 cells was prevented by the expression of flagellin in the parental and *ΔflgM* strains (fig. S3C). Furthermore, the expression of high levels of flagellin by the *ΔflgM* bacteria was associated with more persistent colonization in the intestinal lumen (Fig. 2F), suggesting that flagellin can promote host/bacteria symbiosis through prevention of a pro-inflammatory response. This is in contrast with flagellin expression in the context of intestinal infection by *Salmonella typhimurium (16)*, which induces high levels of IFN*γ* expression by lymphoid cells *(17, 18)* and leads to significant intestinal damage *(19)*.

Even though flagellin is expressed as a cytoplasmic protein in *Villin*^rtTA^ x c*fliC* mice, flagellin-induced activation of epithelial cells and the expansion of Tregs was dependent on TLR5 (Fig. 1D and 2C, fig. S2B), and only marginally dependent on the NLRC4-activated pathway that involves the inflammasome and Caspase-1 (fig. S3D). We therefore investigated whether the production of cytokines by epithelial cells expressing flagellin was cell autonomous. Intestinal organoids were grown from intestinal crypts isolated from *Villin*^rtTA^ x c*fliC* mice, and treated with dox to induce the expression of flagellin (Fig. 2G). Remarkably, the expression pattern of cytokines induced by flagellin in isolated epithelial cells was similar to that induced *in vivo*. Such cell-autonomous activation of epithelial cells was phenocopied by exogenous provision of flagellin (Fig. 2H), suggesting that cytoplasmic flagellin expressed in *Villin*^rtTA^ x c*fliC* mice is released by exosomes or dying cells, with subsequent recognition by TLR5.

We next reasoned that recognition of flagellin by epithelial cells, and the subsequent induction of an anti-inflammatory program in the intestine, would make *Villin*^rtTA^ x c*fliC* mice more resistant to pathological inflammation. To test this hypothesis, dox-treated *Villin*^rtTA^ x c*fliC* mice were fed dextran sodium sulfate (DSS) in the drinking water, a treatment that induces mucus erosion and epithelial injury that results in pathological inflammation in the colon and ileum *(20)*. In contrast to our expectation, expression of flagellin induced a more severe pathology (Fig. 3A) characterized by the expansion of Th17 cells and variable expansion of Tregs (Fig. 3B, fig. S4A). These observations are in accordance with previous data showing that recognition of flagellin, in the context of infection and injury, induces a pro-inflammatory response *(6, 16, 17)*. In this context, the expression by epithelial cells of transcripts for type 2 inducer cytokines, as well as of *Tgfb1* and *Aldh1a2*, was blocked, whereas the expression of *Il1b* was maintained, and the expression of *Il18* was increased (Fig. 3C). Moreover, high levels of IL-18 protein were produced in organoid cultures of *Villin*^rtTA^ x c*fliC* intestinal crypts (Fig. 3D). Here, the expansion of Th17 cells and the expression of *Il1b* and *Il18*, the gene product of which are processed by the inflammasome *(21)*, were independent of TLR5 (Fig. 3C, fig. S4B). Neutralization of IL-18, but not of IL-1*β*, attenuated the severity of intestinal pathology (Fig. 3E) and the expansion of Th17 cells (Fig. 3F) that expressed the highest level of IL-18R*α* during DSS challenge (Fig. 3G).

**Fig. 3.**
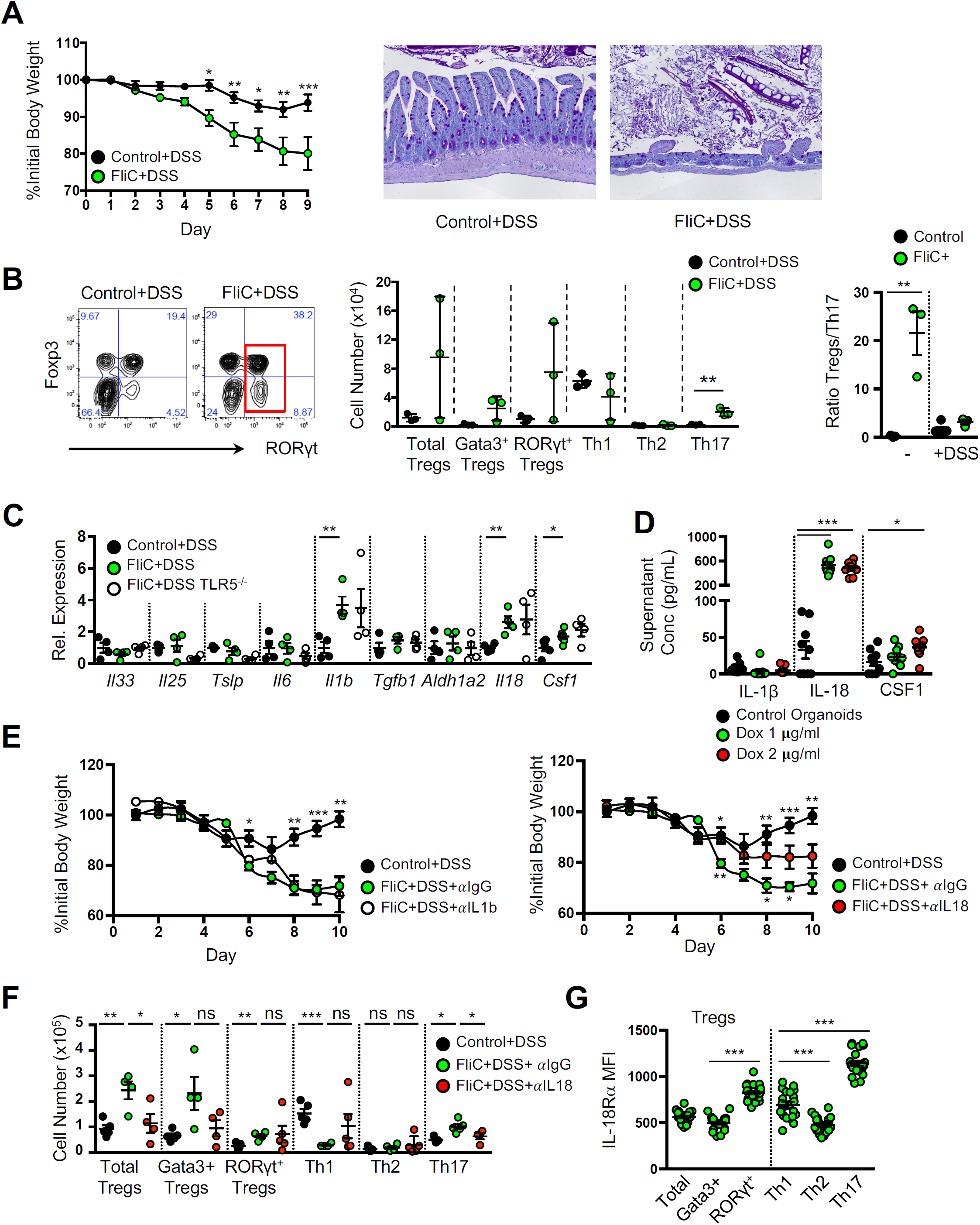
Flagellin-activated epithelial cells induce a pro-inflammatory response in the context of injury. (**A** to **C**) Littermate control, *Villin*^rtTA^ x c*fliC*, and *Villin*^rtTA^ x c*fliC* x *TLR5 KO* mice were treated with dox for 6 days, then with DSS for 7 days. (A) Progression of body weight after DSS treatment, and histopathology 7 days after initiation of DSS. (B) Expression of Foxp3 and ROR*γ*t by T cells (left), cell numbers of Tregs and T helper cells in the small intestine lamina propria of DSS-treated mice (middle), and the ratio of Tregs to Th17 cells in the context of steady state or DSS-induced colitis. (**C**) Gene expression in IECs from the ileum. (**D**) Expression of IL-1*β*, IL-18 and CSF1 by intestinal organoids derived from from *Villin*^rtTA^ x c*fliC* mice. Dox was added 2 days after initiation of the cultures and cytokines measured by ELISA after 7 days. (**E**) Littermate control and *Villin*^rtTA^ x c*fliC* mice were treated with DSS for 5 days, then with dox and either isotype, anti-IL-1*β* or anti-IL-18 neutralizing antibody for 6 days. Progression of body weight during DSS treatment. Asterisks above represent significance between Control+DSS vs. FliC+DSS+αIgG and below represent FliC+DSS+αIgG vs. FliC+DSS+αIL18. (**F**) Cell numbers of Tregs and T helper cells in the small intestine lamina propria of mice treated with anti-IL-18 antibody as in (E). (**G**) Expression of IL-18R*α* by Tregs and T helper cells form the small intestine lamina propria in dox-treated *Villin*^rtTA^ x c*fliC* mice 10 days after initiation of DSS treatment. Error bars show ± SEM, ns means not significant, *P<0.05, **P<0.01, ***P<0.001, as calculated by Student’s *t*-test.

Uematsu *et al.* demonstrated that the detection of flagellin in the context of *Salmonella typhimurium* infection leads to the expansion of Th1 and Th17 cells induced by CD11c^+^ lamina propria cells *(16, 22)*. In dox-treated *Villin*^rtTA^ x c*fliC* mice, the expression of flagellin efficiently induced the recruitment of both CD103^+^ dendritic cells (DCs) and F4/80^+^ macrophages into the intestinal lamina propria, independently of DSS treatment (Fig. 4A and B, fig. S5A). However, the total expression of *Il1b* and *Il18* by intestinal DCs and macrophages from dox-treated *Villin*^rtTA^ x c*fliC* mice was increased in the context of DSS treatment (Fig. 4C), suggesting that myeloid cells were a significant source of pathogenic IL-18. Thus, as epithelial cells in dox-treated *Villin*^rtTA^ x c*fliC* mice expressed high levels of transcripts for the monocyte and macrophage stimulating factor CSF1 (Fig. 2 and 3C), we assessed whether neutralization of CSF1R could prevent the induction of severe intestinal pathology. As expected, CSF1R neutralization blocked the recruitment of both DCs and macrophages (Fig. 4D) and as a consequence, prevented the expansion of Th17 cells (Fig. 4E) and attenuated pathology (Fig. 4F). In accordance with a central role played by epithelial cells in the recruitment of myeloid cells and activation of Th17 cells, organoids grown from intestinal crypts isolated from *Villin*^rtTA^ x c*fliC* mice, or their culture supernatant, were injected into the peritoneum. Organoids treated with dox before transfer induced massive recruitment of macrophages (Fig. S5B) while their culture supernatant induced the activation of Th17 cells (Fig. S5C).

**Fig. 4.**
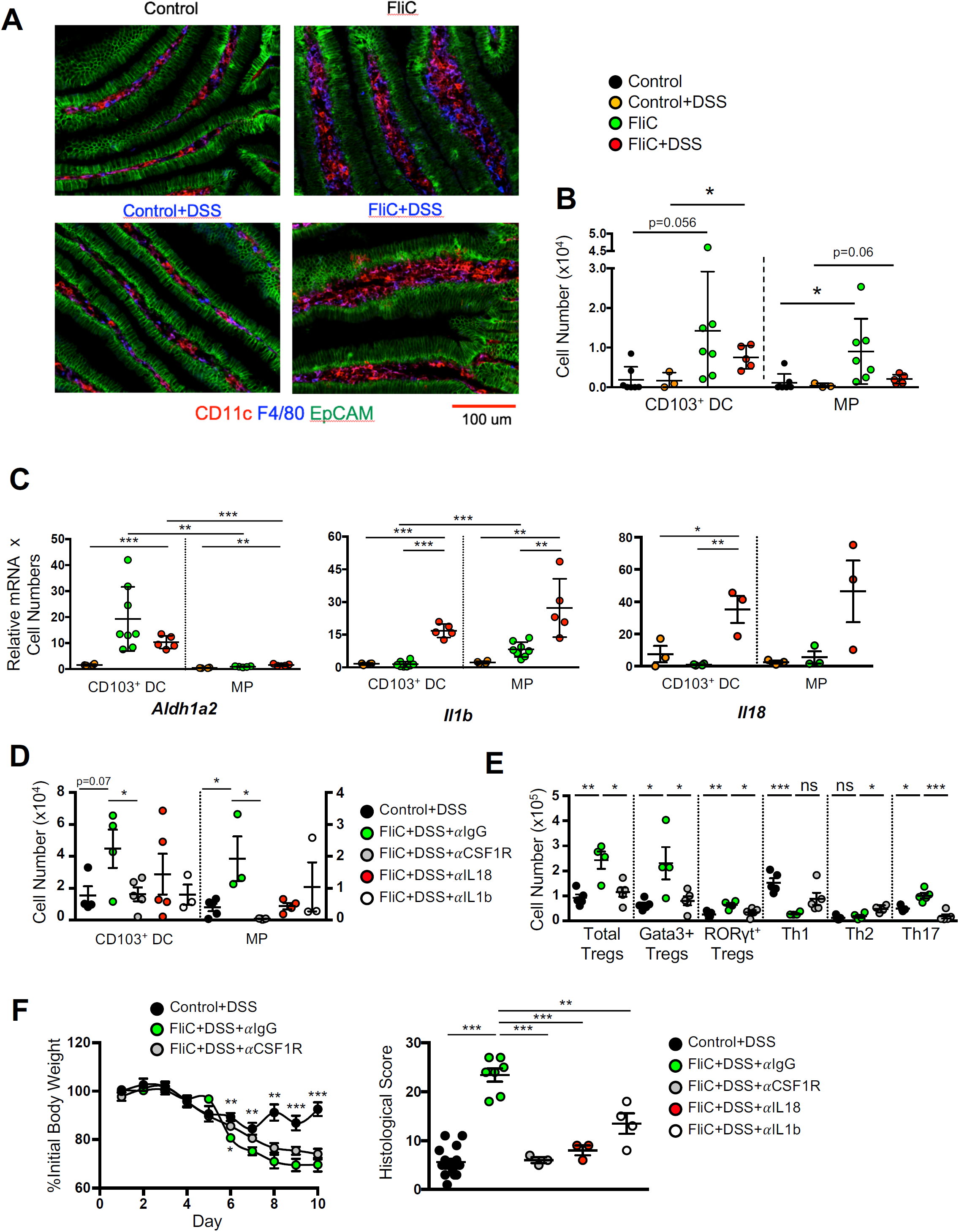
Pro-inflammatory myeloid cells are recruited by epithelial cells in the context of intestinal injury. (**A** to **C**) Dox-treated littermate control and *Villin*^rtTA^ x c*fliC* mice were exposed to DSS for 7 days. Immunofluorescence staining of DCs and macrophages in the ileum (A), cell numbers of CD103^+^ DCs and F4/80^+^ macrophages in the small intestine lamina propria (B) and total expression of transcripts for Aldh1a2, IL-1*β* and IL-18 in the populations of CD103^+^ DCs and macrophages (C). (**D** to **F**) Littermate control and *Villin*^rtTA^ x c*fliC* mice were treated with DSS for 5 days, then with dox and either isotype or anti-IL-1*β*, anti-IL-18 or anti-CSF1R neutralizing antibody for 6 days. The number in the small intestine of CD103^+^ DCs and F4/80^+^ macrophages (D), of Tregs and T helper cells (E) and the progression of body weight and histological scores after initiation of DSS treatment (F). Asterisks above represent significance between Control+DSS vs. FliC+DSS+αIgG and below represent FliC+DSS+αIgG vs. FliC+DSS+αCSF1R. Error bars show ± SEM, ns means not significant, *P<0.05, **P<0.01, ***P<0.001, as calculated by Student’s *t*-test.

These data demonstrate that, at the steady state, the expression and recognition of flagellin by epithelial cells lead to an anti-inflammatory response mediated by IL-33-dependent Tregs. In contrast, in the context of tissue injury, expression of flagellin lead to a pro-inflammatory and pathogenic response mediated by myeloid cells, IL-18 and Th17 cells. Thus, the recognition of flagellin by the innate immune system does not determine the nature of the immune response (originally predicted to be primarily pro-inflammatory)*(1, 2)*. Rather, the recognition of flagellin activates the innate immune system into a way determined by the physiological state of the tissue. In the absence of infection and injury, both the host and its flagellin-expressing microbiota benefit from an anti-inflammatory response. In contrast, infection and injury is countered by a pro-inflammatory response to flagellin and other immunogens. We have previously suggested a similar concept for the generation of microbiota-induced Th17 cells and type 3 (ROR*γ*t^+^) Tregs in the intestine *(12)*. The ratio of Th17 versus Tregs cells, i.e. the pro-inflammatory tone of the anti-microbial response, is determined by the level of RA converted from dietary vitamin A by intestinal cells that may reflect the physiologic state of the tissue. More generally, individual microbes may be defined as symbionts or pathogens, not intrinsically, but rather in the context of their interaction with the host’s tissues. This distinction is important for our understanding of the immunogenic or tolerogenic properties of microbes and microbial compounds.

## Acknowledgements

We thank all the members of the Microenvironment & Immunity Unit, as well as from the Stroma, Inflammation & Tissue Repair Unit, for support and discussion and the members of Gnotobiology Platform of the Institut Pasteur for technical support with GF mice.

## Author Contributions

G.E. conceived the project with critical input from I.G.B.; J.H.P and S.C. initiated the project and performed most experiments; G.N. and P.S. coordinated and performed experiments with organoids; G.S., A.G., B.C. and N.B. performed monoloconization experiments with *E. coli* mutants; F.D. and F.L. generated the transgenic mice; R.S. provided the Villin-rtTA mice; M.B. coordinated experiments on monocolonized germfree mice; G.E., J.H.P. and S.C. wrote the manuscript; all authors corrected the manuscript.

## Data and materials availability

All data is available in the manuscript or the supplementary materials.

## Supplementary Materials

### Materials and Methods

#### Mice

BAC-transgenic transgenic *cfliC* mice were generated as described previously *(23)*. The *fliC* coding sequence derived from *Salmonella typhimurium* was codon-optimized for the mouse genome and inserted into a “neutral” bacterial artificial chromosome (BAC) (Invitrogen) devoid of genes, carrying 200kb of the genomic region targeted in the transgenic mouse line LC1 *(24)*. Expression of the *fliC* coding sequence was controlled by a dox-inducible *tetO*_*7*_ promoter, and followed by an *Ires-EGFP* sequence. In *sfliC* mice, the *fliC* coding sequence was preceded by a sequence coding for the signal peptide of IL-7. All mice were backcrossed to C57Bl/6 for at least 5 generations. In all experiments, wild-type, *Villin*^rtTA^ or *cfliC* single positive transgenic littermates were used as controls. C57BL/6 mice were purchased from Charles River and TLR5-deficient mice were obtained from Shizuo Akira. All mice were kept in specific pathogen-free conditions. Germfree mice were generated from C57BL/6 mice and maintained at the Gnotobiology Platform of the Institut Pasteur. All animal experiments were approved by the committee on animal experimentation of the Institut Pasteur and by the French Ministry for Higher Education, Research and Innovation.

#### Mouse treatments

For the analysis of flagellin response in C57BL/6 mice, 5 μg of ultrapure flagellin from *Salmonella typhimurium* (Invivogen) was injected intraperitoneally daily for 6 days. For flagellin expression in intestinal epithelial cells, *Villin*^rtTA^ *x cfliC* mice were fed 1 mg/ml doxycycline (dox) in their drinking water for 6 days. For epithelial tissue injury, 6 days of dox treatment were followed by 7 days of 2.5% DSS (MP Biochemicals) in their drinking water. For TLR5 or IL-33 neutralization, 50 μg of anti-TLR5-IgG (Invivogen) or 25 μg of anti-IL-33 antibody (R&D) were administered intraperitoneally daily during 6 days of dox treatment. Antibody neutralization of IL-1*β* (B122), IL-18 (YIGF74-1G7) or CSF1R (AFS98; 8mg/kg; Bioxcell) was performed after 5 days of DSS while dox was administered in the drinking water for 6 days. For inhibition of the retinoic acid receptor, 330 μg of the pan-retinoic acid receptor inverse agonist BMS493 (R&D) was administered intraperitoneally daily during the 6 days of dox treatment. For inhibition of caspase-1, 100 μg of Ac-YVAD-cmk (Invivogen) was administered intraperitoneally daily during dox or DSS treatment. For the mono-colonization of germfree mice with *E. coli* strain LF82, *ΔflgM* or *ΔfliC*, bacteria were grown overnight in 200 mL of LB at 37°C in Luria-Bertani (LB) broth containing ampicillin (100 μg/ml), erythromycin (20 μg/ml) for wildtype, and kanamycin (50 μg/ml) for mutant strains. Bacterial suspensions with an OD_620nm_ of 2.0 were placed in the water bottles of germ-free C57Bl/6 mice which had been placed in isolated ventilated caging system (Isocage) that prevents exogenous bacterial contamination. Two or 8 weeks later, mice were euthanized, and their organs were collected for downstream analysis.

#### Cell preparation and culture

Small intestinal and colonic lamina propria cells were isolated as described previously *(11, 12)*. Briefly, Peyer’s Patches were removed, and whole small intestine and colon were opened longitudinally, cut into pieces and incubated for 30 minutes in 30mM EDTA, washed extensively, and incubated for 30 min at least twice in 50-75 μg/ml liberase TL (Roche). These preparations were then pushed through a 100μm filter to generate single cell suspensions. Cells were separated by a 40/80% (w/v) Percoll (GE Healthcare) density gradient and washed prior to staining for flow cytometry analysis. For isolation of epithelial cells of small intestine, Peyer’s patches were removed, and the proximal 1/3 (duodenum) or distal 1/3 (ileum) of small intestine were opened longitudinally, cut into pieces, and incubated for 30 min in DMEM containing 5 mM EDTA with shaking (180 rpm) at 37C. Cells were filtered through a 100 μm filter, and pelleted. RNA was prepared using RNeasy mini kit (Qiagen). For analysis of cytokine production in T cells, cells were cultured for 3 days in round-bottomed 96-well plates coated with anti-CD3 (10 μg/ml) in RPMI 1640 (Invitrogen) containing soluble anti-CD28 (1 μg/ml) and 10% FCS, 25 mM HEPES, 0.05 mM 2-mercaptoethanol, and 100 U/ml of penicillin and streptomycin. Cytokine levels in culture supernatants were analysed by ELISA (eBioscience). For organoid formation, intestinal crypts were extracted as previously described *(25)*. Organoids were treated with ultrapure flagellin or dox 2 days after crypt incubation, and further incubated for 5 days.

#### Flow cytometry

Cells were pre-incubated with LIVE/DEAD Fixable Blue Dead Cell Stain Kit for 30 minutes, Fc-Block for 15 minutes, and stained for 20 minutes with the following antibodies: Brilliant Violet 785-conjugated anti-CD3 (17A2; BioLegend) or PerCP-Cy-5.5-conjugated anti-CD3 (17A2), Horizon V500-conjugated anti-CD4 (RM4-5; BD), FITC or V500-conjugated anti-CD45.2 (104; BD), Brilliant Violet 650-conjugated anti-CD19 (6D5, Biolegend), eFluor450-conjugated anti-MHC II (M5/114.15.2), eFluor660-conjugated anti-CD11c (N418), PE-conjugated anti-CD103 (2E7), PE-Cy7-conjugated anti-CD11b (M1/70), Alexa Fluor 488-conjugated anti-F4/80 (BM8), biotin-CD218a (REA947; Miltenyi Biotec). For intracellular staining, cells were fixed and permeabilized with a commercially available fixation/permeabilization buffer (eBioscience). Intracellular staining was performed with the following antibodies: Pacific Blue-conjugated anti-Helios (22F6; BioLegend), PE-conjugated anti-RORγt (AFKJS-9), PerCP-Cy5.5-conjugated anti-Foxp3 (FJK-16s), eFluor660-conjugated anti-Gata3 (TWAJ), PE-Cy7-conjugated anti-T-bet (4B10). All antibodies were from eBioscience unless stated otherwise. For *in vivo* organoid recruitment studies involving Th cells, intestinal organoids were grown as above and spent supernatant (500uL) from *Villin*^rtTA^ *x cfliC* treated with or without dox was injected into the peritoneum. A peritoneal lavage was performed after 24 hours and Th17 (eGFP^+^), iTreg (eGFP^+^RFP^+^) and Gata3^+^ (eGFP^-^RFP^+^) Tregs were sorted (CD3^+^CD4^+^CD45^+^Live) on a FACS Aria III directly into RLT buffer and RNA extracted with the Qiagen RNAeasy Micro kit.

#### Immunofluorescence histology

Tissues were fixed and stained as previously described *(26)*. In brief, tissues were washed and fixed overnight at 4°C in 4% paraformaldehyde (Santa Cruz). The samples were then washed for 2 days in PBS, incubated in a solution of 30% sucrose (Sigma-Aldrich) in PBS, and embedded in OCT. Frozen blocks were cut at 8-μm thickness and sections collected onto Superfrost Plus slides (VWR). For staining, sections were first hydrated in PBS-TS (PBS containing 0.1% Triton X-100 and 1% bovine serum; Sigma-Aldrich) and blocked with 10% bovine serum in PBS-TS for 1 hour at room temperature. Sections were incubated with primary antibody or conjugated antibody in PBS-TS overnight at 4°C, washed with PBS-TS, incubated with secondary antibody for 1 hour at room temperature if necessary. Sections were incubated with DAPI for 5 min at room temperature, washed with PBS-TS, and mounted with Fluoromount-G. Slides were examined under a fluorescence microscope (AxioImager M1; Carl Zeiss, Inc.). Sections were stained with the following antibodies; unconjugated polyclonal anti-GFP (rabbit), unconjugated anti-E-cadherin (ECCD-2), Alexa Fluor 488-conjugated anti-rabbit IgG secondary antibody, Alexa Fluor 555-conjugated anti-rat IgG secondary antibody were from Invitrogen. APC-conjugated anti-EpCAM (G8.8), PE-conjugated anti-CD11c (N418), APC-conjugated anti-F4/80 (BM8) were from eBioscience. For *in vivo* organoid retrieval studies to track macrophage recruitment, intestinal organoids from *Villin*^rtTA^ *x cfliC* were grown as above with and without dox, recovered without dissociation and injected into anesthetized mice with a 22g needle to prevent shearing. After 4 hours, a peritoneal lavage was performed, peritoneal contents spun at 500g and processed for immunofluorescence as above and imaged on a Leica SP8 laser scanning confocal in 3D.

#### Histopathology

Ileum segments were fixed overnight at 4°C in 4% paraformaldehyde and dehydrated through an ethanol gradient before embedding in paraffin. Tissue blocks were sectioned at 5 micron thickness and stained with Periodic Acid-Schiff with counterstaining to hemotoxylin. Slides were scanned at 20X with a Axio Scan Z1 (Zeiss) and blindly scored for disease activity as based on the following parameters: Inflammatory infiltrate severity (1-4) and extent (1-3), epithelial cell damage including erosion (1-4), ulceration (3-5), irregular crypts (4-5), villous blunting (3-5) and goblet cell depletion (1-4), with 5 being the worst score.

#### Cell death and proliferation

Mice were injected with 50 μl of 10 mM 5-ethynyl-2’-deoxyuridine (EdU, Invitrogen) solution intraperitoneally 1 day before analysis. Incorporation of EdU into proliferating cells was detected by histology with the Click-iT EdU Alexa Fluor 555 Imaging Kit (Invitrogen), according to the manufacturer’s protocol.

#### Quantitative PCR

To isolate RNA from FACS-sorted cells, cells were directly sorted into RLT buffer (Qiagen) supplemented with *β*2-mercaptoethanol. Total RNA was prepared using RNeasy micro kit (Qiagen) according to the manufacturer’s instructions. To prepare RNA from whole tissues, tissues from terminal ileum and proximal colon were homogenized in Trizol using gentleMACS Octo Dissociator (Miltenyi). Chloroform was added, and aqueous phase was separated by centrifugation. RNA is precipitated from the aqueous layer with isopropanol, and further purified using RNeasy mini kit (Qiagen). Concentration and integrity were assessed using the Bioanalyzer (Agilent). cDNA was synthesized by reverse transcription using Superscript IV (Invitrogen). Detection of specific gene expression was performed using RT2 qPCR primers (Qiagen) and SYBR green (Biorad). Gene expression was normalized to *Gapdh, Hsp90* and *Hprt* in each sample.

#### Statistical analysis

P-values were calculated with unpaired, two-tailed *t* test or Mann-Whitney test using GraphPad PRISM 6 software.

### Supplementary Figure legends

**Fig. S1. Apoptosis or proliferation of IEC was not increased upon flagellin expression in *Villin***^**rtTA**^ **x c*fliC* mice.** Representative pictures of Tunel assay and EdU immunostaining in IECs of control and *Villin*^rtTA^ x c*fliC* mice after 6 days of doxycycline (dox) treatment in drinking water.

**Fig. S2. Expansion of type 2 immune responses in *Villin***^**rtTA**^ **x c*fliC* mice.**

(**A**) Frequencies of Th17 cells, and frequencies and cell numbers of ILC3s in small intestine lamina propria of littermate control and *Villin*^rtTA^ x c*fliC* mice treated with recombinant flagellin intraperitoneally daily for 6 days. (**B**) Frequencies of Tregs in the small intestines of littermate control and *Villin*^rtTA^ x c*fliC* mice treated with either isotype or anti-TLR5 neutralizing antibody. (**C**) Cell numbers and frequencies of Th2, Th17, ILC2 and ILC3 in the small intestines of control and *Villin*^rtTA^ x c*fliC* mice treated with dox. (**D**) Frequencies of Tregs in the small intestines of littermate control and *Villin*^rtTA^ x *sfliC* mice treated with dox. Error bars show ± SEM, *P<0.05, **P<0.01, as calculated by Student’s *t*-test.

**Fig. S3. Flagellin induces epithelial cells to promote type 2 Tregs.** (**A**) Frequencies of Tregs in control and *Villin*^rtTA^ x c*fliC* mice treated with dox and either DMSO or retinoic acid receptor (RAR) inhibitor intraperitoneally daily for 6 days. (**B**) Cell numbers of Tregs in control and *Villin*^rtTA^ x c*fliC* mice treated with dox and either isotype or anti-IL-33 neutralizing antibody. (**C**) Relative cell numbers of Tregs and T helper cells in the small intestines of germ-free mice colonized with wild-type, *ΔflgM* or *ΔfliC E.coli* LF82 for 2 weeks. (**D**) Frequencies and cell numbers of Tregs in control and *Villin*^rtTA^ x c*fliC* mice treated with dox and either PBS or Caspase-1 inhibitor intraperitoneally daily for 6 days. Error bars show ± SEM, ns means not significant, *P<0.05, **P<0.01, ***P<0.001, as calculated by Student’s *t*-test.

**Fig. S4. The expansion of Th17 cells is independent of TLR5 during DSS-induced colitis.**

(**A**) Littermate control and *Villin*^rtTA^ x c*fliC* were treated with dox for 6 days, then with DSS for 7 days, and frequencies of Tregs and T helper cells measured in the small intestine. (**B**) Littermate control, *Villin*^rtTA^ x c*fliC*, and *Villin*^rtTA^ x c*fliC* x *TLR5 KO* mice were treated as in (A), and cell numbers and frequencies of Tregs and T helper cells measured in the small intestines. Error bars show ± SEM, ns means not significant, *P<0.05, **P<0.01, ***P<0.001, as calculated by Student’s *t*-test.

**Fig. S5. Pro-inflammatory myeloid cells are recruited by epithelial cells in the context of intestinal injury.** (**A**) Dox-treated littermate control and *Villin*^rtTA^ x c*fliC* mice were exposed to DSS for 7 days. Immunofluorescence staining of DCs and macrophages in the ileum. (**B**) Intestinal organoids derived from *Villin*^rtTA^ x c*fliC* mice were grown for 5 days with and without dox and transferred into the peritoneum of C57BL/6 mice. After 4 hours, organoids were isolated and analysed by immunofluorescence for the recruitment of F4/80^+^ macrophages. Images shown represent a 3D stack maximally projected to show all planes. (**C**) Supernatant from intestinal organoids was injected into the peritoneum of *Foxp3-RFP x RORgt-EGFP* mice for 24 hours whereby recruitment of Tregs and Th17 cells was assessed by FACS sorting followed by qPCR of *Il-17a* to assess the activation state of Th17 cells. Error bars show ± SEM, ns means not significant, *P<0.05, **P<0.01, ***P<0.001, as calculated by Student’s *t*-test.

