## Supplementary figures and images for "Innate immune recognition of a bacterial MAMP leads to conditional activation of pro- or anti-inflammatory responses"

### Supplemental Figures

#### Figure S1

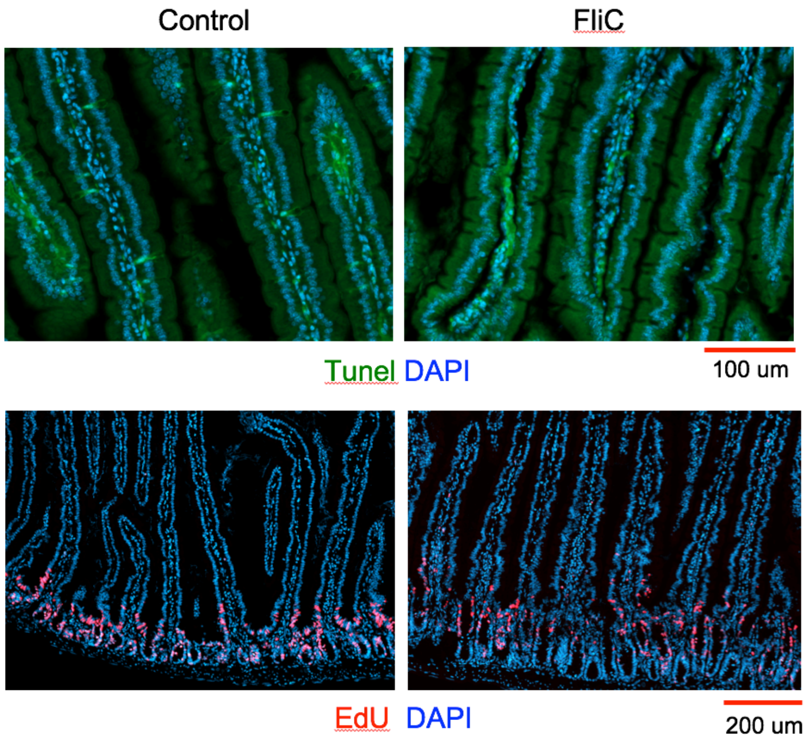

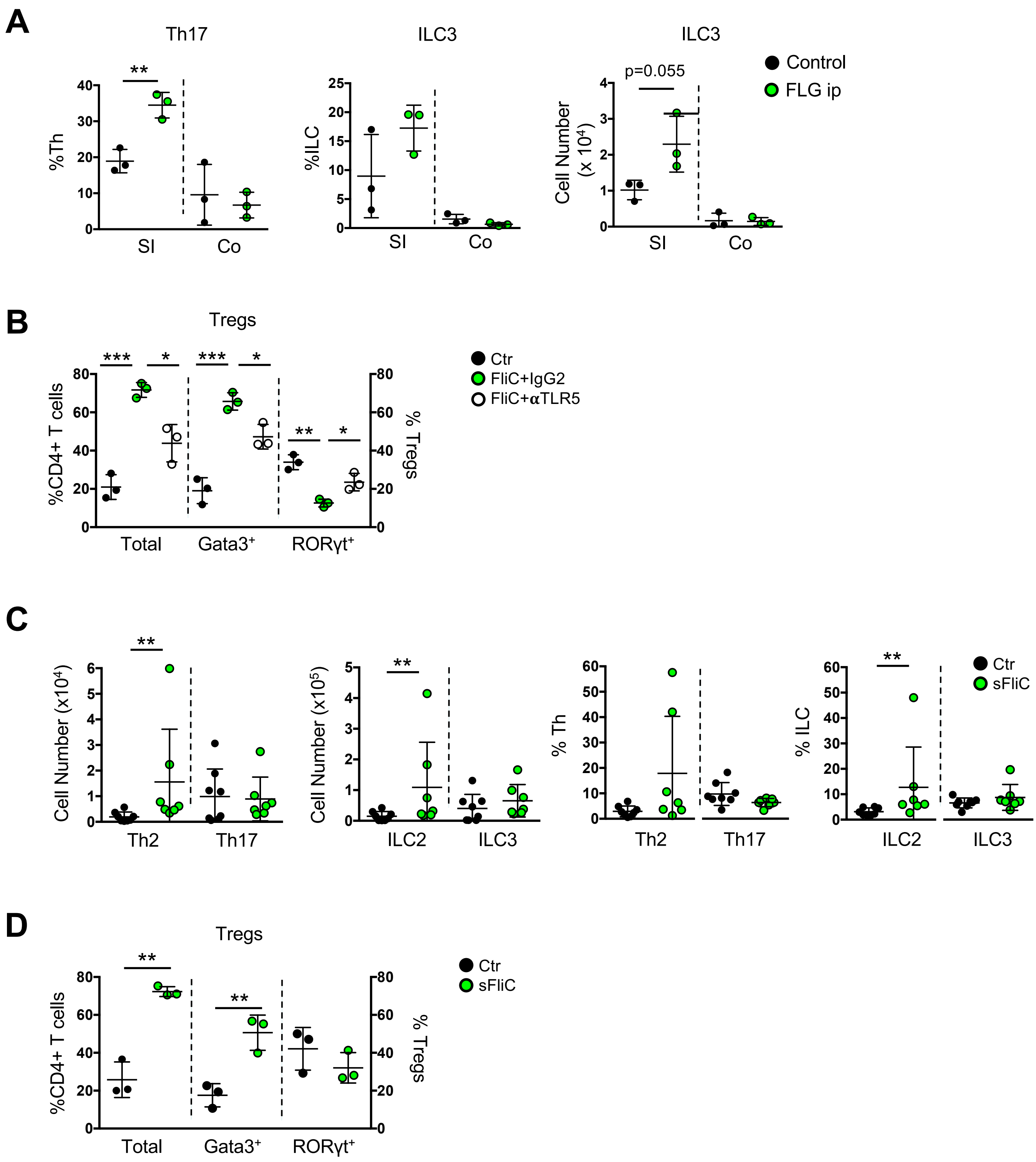

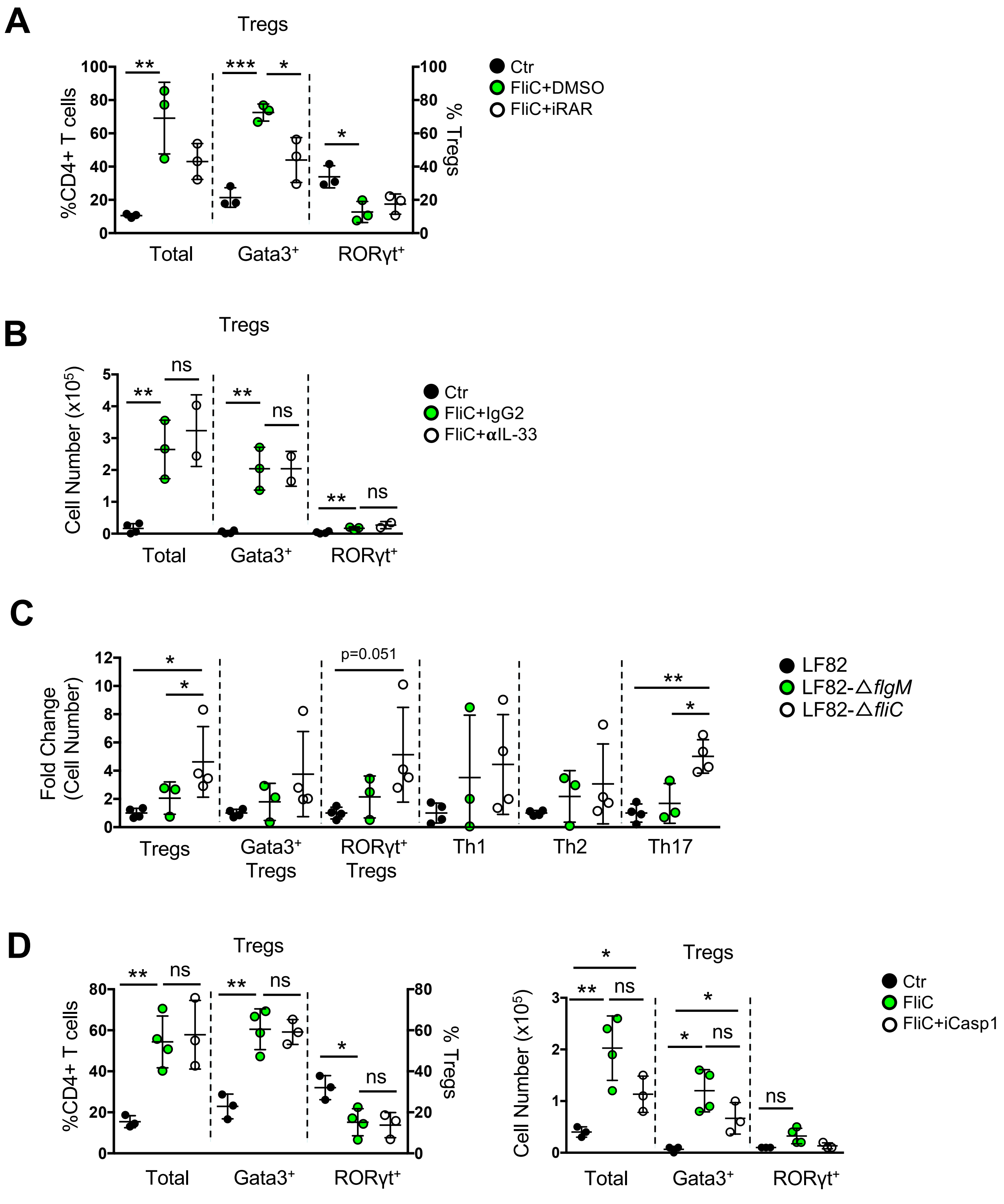

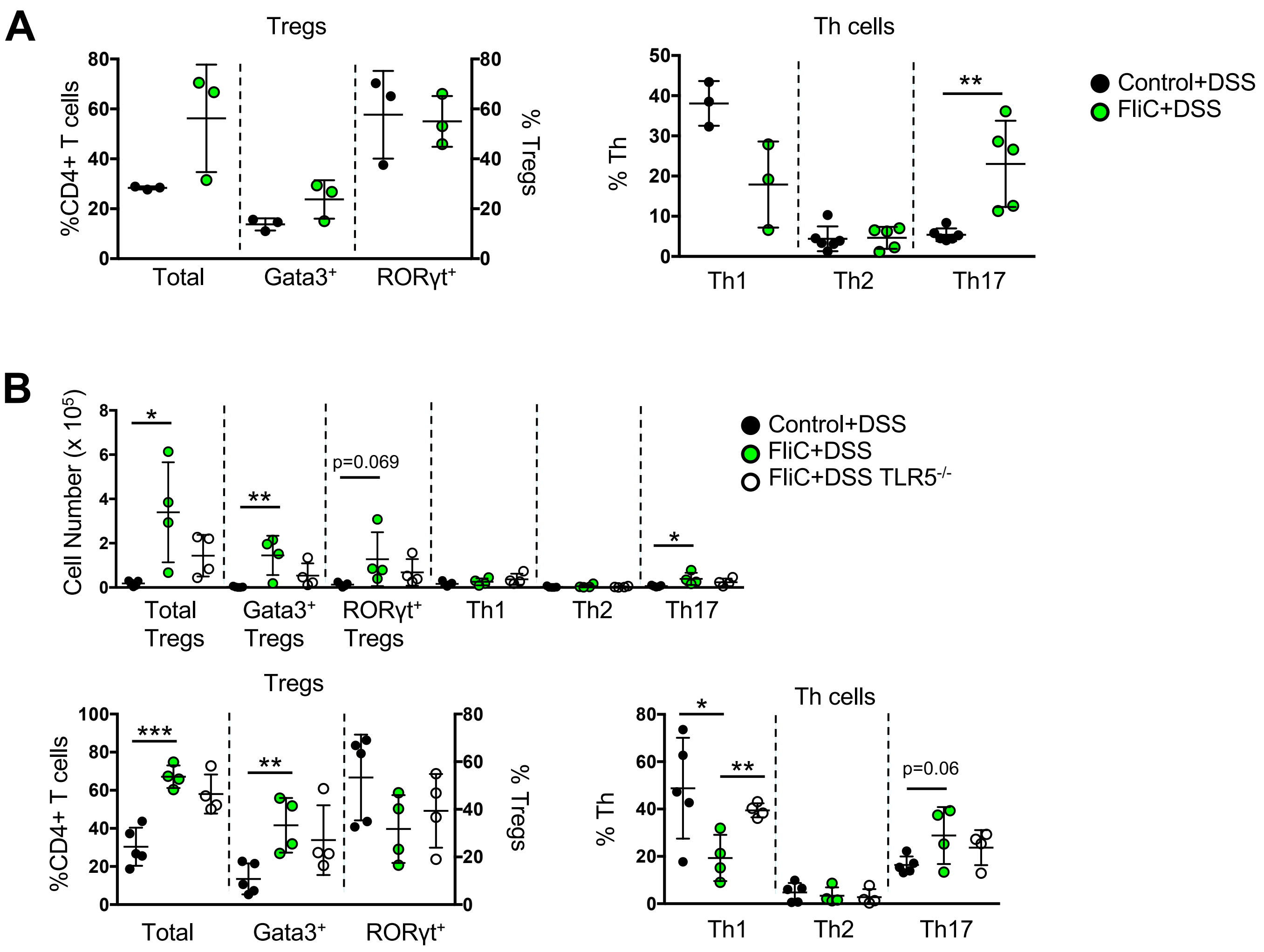

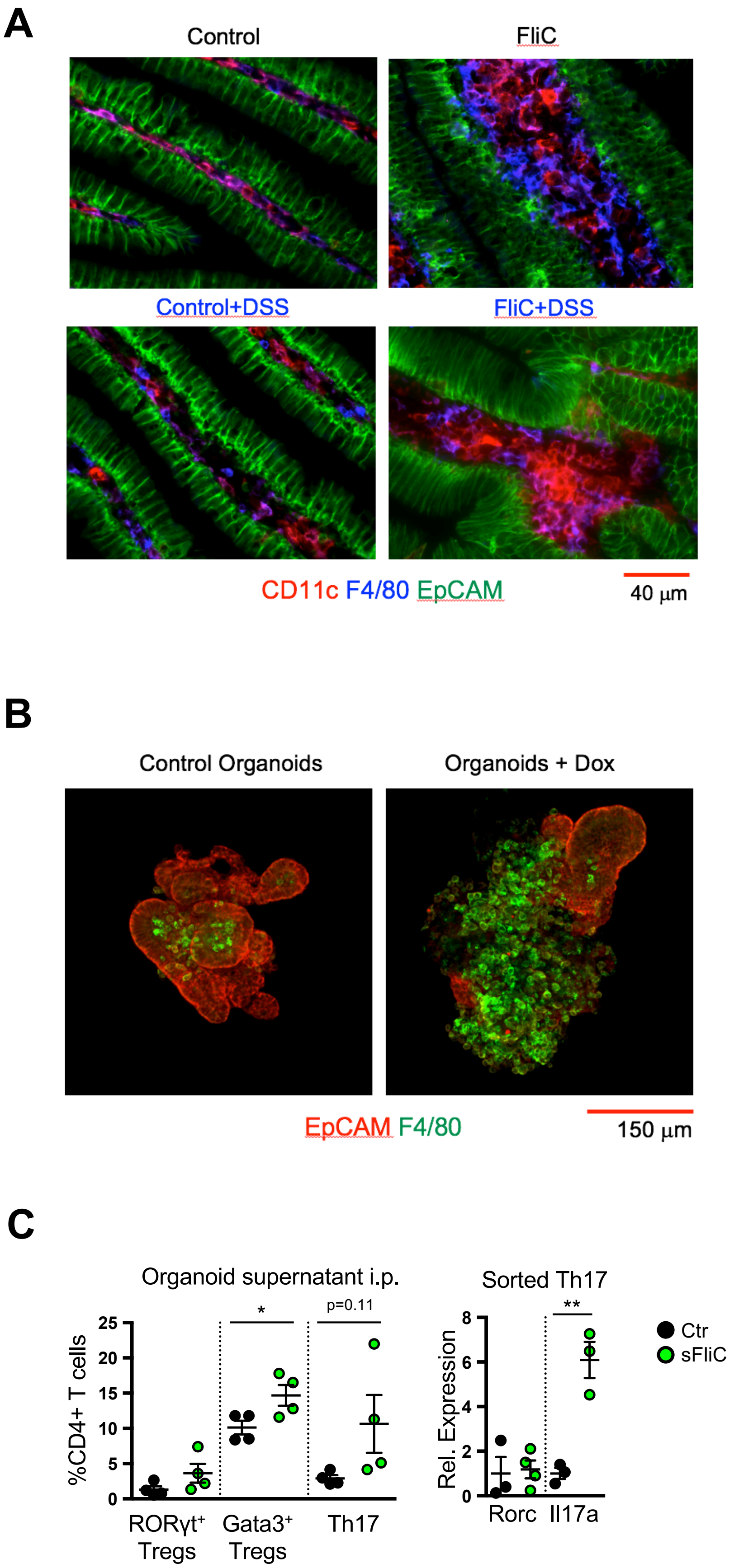
